# The cryo-EM structure of vesivirus 2117 highlights functional variations in entry pathways for viruses in different clades of the Vesivirus genus

**DOI:** 10.1101/2021.02.05.429895

**Authors:** Hazel Sutherland, Michaela J. Conley, Edward Emmott, James Streetley, Ian G. Goodfellow, David Bhella

**Affiliations:** Medical Research Council – University of Glasgow Centre for Virus Research, Sir Michael Stoker Building, Garscube Campus, 464 Bearsden Road, Glasgow G61 1QH, United Kingdom; Scottish Centre for Macromolecular Imaging, Sir Michael Stoker Building, Garscube Campus, 464 Bearsden Road, Glasgow G61 1QH, United Kingdom; Division of Virology, Department of Pathology, University of Cambridge, Addenbrooke’s Hospital, Hills Road, Cambridge CB2 2QQ, United Kingdom; Centre for Proteome Research, Institute of Systems, Molecular & Integrative Biology, University of Liverpool, Liverpool, L69 7ZB, United Kingdom

## Abstract

Vesivirus 2117 is an adventitious agent that has been responsible for lost productivity in biopharmaceutical production following contamination of Chinese hamster ovary cell cultures in commercial bioreactors. A member of the *Caliciviridae*, 2117 is classified within the Vesivirus genus in a clade that includes canine and mink caliciviruses but is distinct from the vesicular exanthema of swine clade, which includes the extensively studied feline calicivirus (FCV). We have used cryogenic electron microscopy (cryo-EM) to determine the structure of the capsid of this small, icosahedral, positive-sense RNA containing virus. We show that the outer face of the dimeric capsomeres, which contains the receptor binding site and major immunodominant epitopes in all caliciviruses studied thus far, is quite different from that of FCV. This is a consequence of a 22 amino-acid insertion in the sequence of the FCV major capsid protein that forms a ‘cantilevered arm’, which plays an important role in both receptor engagement and undergoes structural rearrangements thought to be important for genome delivery to the cytosol. Our data highlight a potentially important difference in the attachment and entry pathways employed by the different clades of the Vesivirus genus.

## Introduction

Vesivirus 2117 is an adventitious agent that was first identified following contamination of bioreactors containing Chinese Hamster Ovary (CHO) cells for biopharmaceutical production at a Genzyme manufacturing plant in Allston, Massachusetts (USA) (1). The virus was believed to have been introduced via a cell culture reagent, therefore the natural host is currently unknown. The incident halted production of two medications used to treat rare, inherited enzyme-deficiency disorders. The standstill in production during decontamination reportedly cost the company $100-300 million and affected around 8000 patients (2).

Vesivirus 2117 is a non-enveloped, single stranded positive sense RNA virus that belongs to the *Caliciviridae*. Caliciviruses are important pathogens that infect both humans and animals. There are eleven genera including the Vesiviruses, Noroviruses, Lagoviruses and Sapoviruses (3). Noroviruses and Sapoviruses are the most clinically significant as they are major causative agents of gastroenteritis in humans (4). Vesiviruses infect animal hosts and cause wide ranging symptoms from respiratory illness to haemorrhagic disease. The best understood Vesivirus is feline calicivirus (FCV), which causes respiratory disease in cats. Virulent systemic forms of FCV have recently emerged, associated with high mortality rates (5). The natural host of vesivirus 2117 is not known, however 2117-like viruses have been isolated from dogs showing symptoms of severe gastroenteritis and phylogenetic analysis places 2117 in the same clade as canine caliciviruses (CaCV) (6, 7). Antibodies to 2117-like viruses have also been detected in humans and cats (8, 9).

Vesivirus genomes are approximately 7.5kb in length and contain three open reading frames (ORFs). ORF1 encodes a large polyprotein that is cleaved into the various non-structural proteins including the helicase, protease and RNA dependent RNA polymerase (10). ORF2 encodes the major capsid protein VP1 while ORF3 encodes the minor capsid protein VP2, which has been shown to be essential for virus replication (11–13). In common with all known caliciviruses, vesivirus 2117 assembles a T=3 icosahedral capsid that is therefore made up of 180 copies of the major capsid protein VP1 (6). There are 3 quasi-equivalent conformations of VP1, denoted as A, B and C. Together they form the capsid asymmetric unit. These conformers form two types of dimeric capsomeres; AB dimers are arranged about the icosahedral five-fold symmetry axes, while CC dimers are located at the icosahedral two-fold axes. VP1 can be divided into three domains: the P (protruding) domain, the S (shell) domain and the NTA (N-terminal arm). The P domain forms the distinctive arch-shaped spikes on the capsid surface giving rise to the characteristic cup-shaped morphology of the virus from which the *Caliciviridae* derive their name (Greek *calyx*). The P domain is further divided into two subdomains: P2, which is distal to the capsid shell and P1 which forms the base of the protruding capsomere. P2 is an insert in P1 and contains the receptor binding site and major immunodominant epitopes on its outermost face. The S domains form the contiguous capsid floor while the NTA extends from the S-domain at the capsid interior stabilising interactions between AB and CC dimeric capsomeres and directing virion assembly (14).

In FCV, the minor capsid protein VP2 has been shown to assemble a funnel-shaped portal-like structure at a unique three-fold symmetry axis following receptor engagement (15). FCV attachment and entry are mediated by feline junctional adhesion molecule A (fJAM-A) an immunoglobulin-like protein found in tight-junctions (16). Receptor engagement leads to substantial conformational rearrangements; rotation and tilting of the P-domains, accompanied by increased flexibility (17, 18). These conformational changes enable formation of the VP2 portal-like assembly. FCV entry proceeds by endocytosis following virion attachment to fJAM-A. We postulate that VP2 mediates endosome escape by inserting into the endosomal membrane and forming a channel through which the viral genome is released into the cytosol.

We have previously noted that the structure of vesivirus 2117 shows substantial morphological differences to other known Vesivirus structures, having a rounded P2 with pronounced horns of density, compared to the flat, extended P2 of FCV and San Miguel sealion virus (6). In FCV the outer face of P2 forms a platform that is able to engage two copies of fJAM-A. This interaction leads to rearrangement of a loop in VP1 at amino-acid residues 436-448 causing it to raise up to meet the receptor. This results in opening of a cleft on the side of P2 into which VP2 binds – anchoring the portal to the P domain. The striking differences that we have previously observed between vesivirus 2117 and FCV indicate a different mode of receptor engagement and VP2 anchor. Here we present the high-resolution structure of the vesivirus 2117 capsid, solved by cryogenic electron microscopy (cryo-EM) and three-dimensional image reconstruction of virus-like particles (VLPs) produced by heterologous expression of the 2117 major capsid protein VP1. We confirm that the outermost face of P2 is topologically divergent from that of FCV and show that the extended loop that we have previously shown to be important for receptor engagement and VP2 stabilisation in FCV, is an insertion that is not present in 2117. These data indicate that the 2117/CaCV clade of Vesiviruses likely exhibit substantial differences in both receptor engagement and VP2 portal assembly.

## Results and Discussion

### Cryo-EM of vesivirus 2117 virus-like particles

To determine the capsid structure of vesivirus 2117, we used cryo-EM to image virus-like particles (VLPs) produced by heterologous baculovirus expression of the VP1 gene in insect cells. Two-thousand micrographs were recorded on a JEOL CryoARM 300 automated transmission cryo-electron microscope equipped with a Direct Electron DE64 detector, at the Scottish Centre for Macromolecular Imaging. Despite considerable numbers of VLPs appearing as distorted or disrupted particles, a dataset of 15,989 particle images was identified that was able to be processed to yield an icosahedral reconstruction at 3.6 Å resolution (Fig 1A, Movie S1, Fig S1). Following sharpening to enhance high-resolution features for model building, P2 showed noisy and discontinuous density. Local resolution analysis also indicated that the P2 domains were not well resolved, showing an estimated resolution of poorer than 4 Å (Fig 1B).

**Figure 1.**
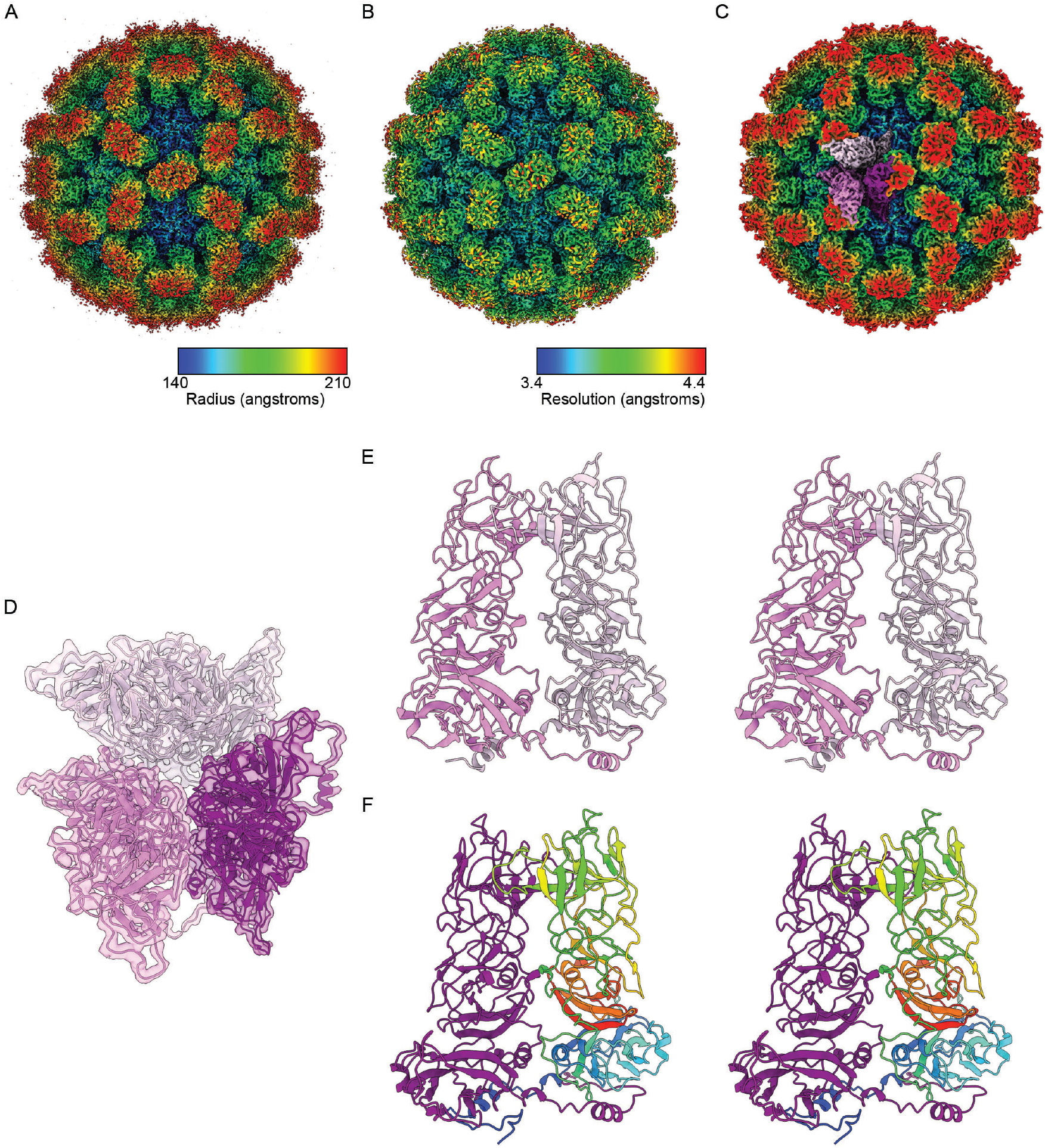
Cryo-electron microscopy reconstructions of the vesivirus 2117 virus-like particle. The structure was determined at 3.6 angstroms resolution. Following sharpening to facilitate visualisation of high-resolution information for model-building, density at the distal tips of capsomeres corresponding to the P2 domain was noisy and difficult to interpret. The sharpened map is shown coloured by radius (A). Local resolution assessment indicated that the P2 region was resolved at poorer than 4 angstroms resolution (B). Selective blurring of features and deep-learning map modification enhanced these features such that a model could be built for 2117 VP1 (C). The modified map is shown coloured by radius and with the locations of quasi-equivalent copies of VP1 in the asymmetric unit highlighted (violet - chain A, thistle - chain B and deep magenta - chain C). A cartoon representation shows the atomic model for the asymmetric unit within a transparent surface to highlight the modified cryo-EM density map (D). Wall-eyed stereo pair images of the AB dimer (E) and CC dimer (F) are shown with rainbow colouring for one copy of VP1 in the CC-dimer to highlight the positions of N-terminus (blue) and C- terminus (red).

### An atomic model of the 2117 asymmetric unit

To assemble an atomic model for VP1 a homology model was generated using the Phyre2 server (19). This was docked to the sharpened cryo-EM density and edited to produce a model for the NTA, S and P1 domains that matched the reconstructed density well. To improve the density for the surface loops in P2, deep learning map modification was applied using DeepEMhancer (20). This gave rise to density that showed improved continuity, allowing modelling of the more challenging regions (Fig 1C). Application of deep-learning map modification is a recently developed and potentially controversial process. Selective blurring of the map yielded similarly interpretable density, however (Fig S2). Furthermore, the presence of quasi-symmetry in the T=3 icosahedral assembly gave rise to features in regions of the map that were not subject to averaging during the reconstruction process but were nonetheless similar, lending confidence to our interpretation.

The resulting VP1 model comprised three chains, one for each quasi-equivalent polypeptide in the asymmetric unit and labelled A, B and C (Fig 1D). The reconstructed density allowed us to model residues Pro10-Gln181 and Thr187-Ala527 for chain A, Pro27-Ala527 for chain B and Ile12-Ala527 for chain C. The model revealed a structure that closely follows the topology of other calicivirus VP1 proteins comprising an extended NTA that lines the internal surface of the capsid, stabilising VP1 dimers and dimer-dimer interactions (Fig 1E,F). Likewise, the S domain exhibits the canonical eight-stranded beta-barrel fold; or beta-jelly roll, found in many small RNA containing spherical viruses. Comparison with the FCV S domain revealed a highly conserved structure, pairwise alignment and comparison gave RMSD values of 1.4 Å for chains A and B and 2.1 Å for chain C (Fig 2A, Movie S2). Similarly, the 2117 VP1 P1 domain shows a consistent architecture when compared to that of FCV (Fig 2B). P2 however shows a highly divergent structure. While P2 domains of FCV and 2117 both comprise a six-stranded antiparallel beta barrel, the surface loops that define the topology of the outer capsid surface show considerable differences. This is perhaps unsurprising, given that this region encodes the receptor binding domains and major immunodominant epitopes for all caliciviruses for which data are available. Such regions in animal viruses are usually characterised by the presence of ornate hyper-variable loops. Comparisons between the P-domains of 2117 and FCV gave RMSD measurements of 6.2 Å for chain A, 6.1 Å for chain B and 7.7 Å for chain C.

**Figure 2.**
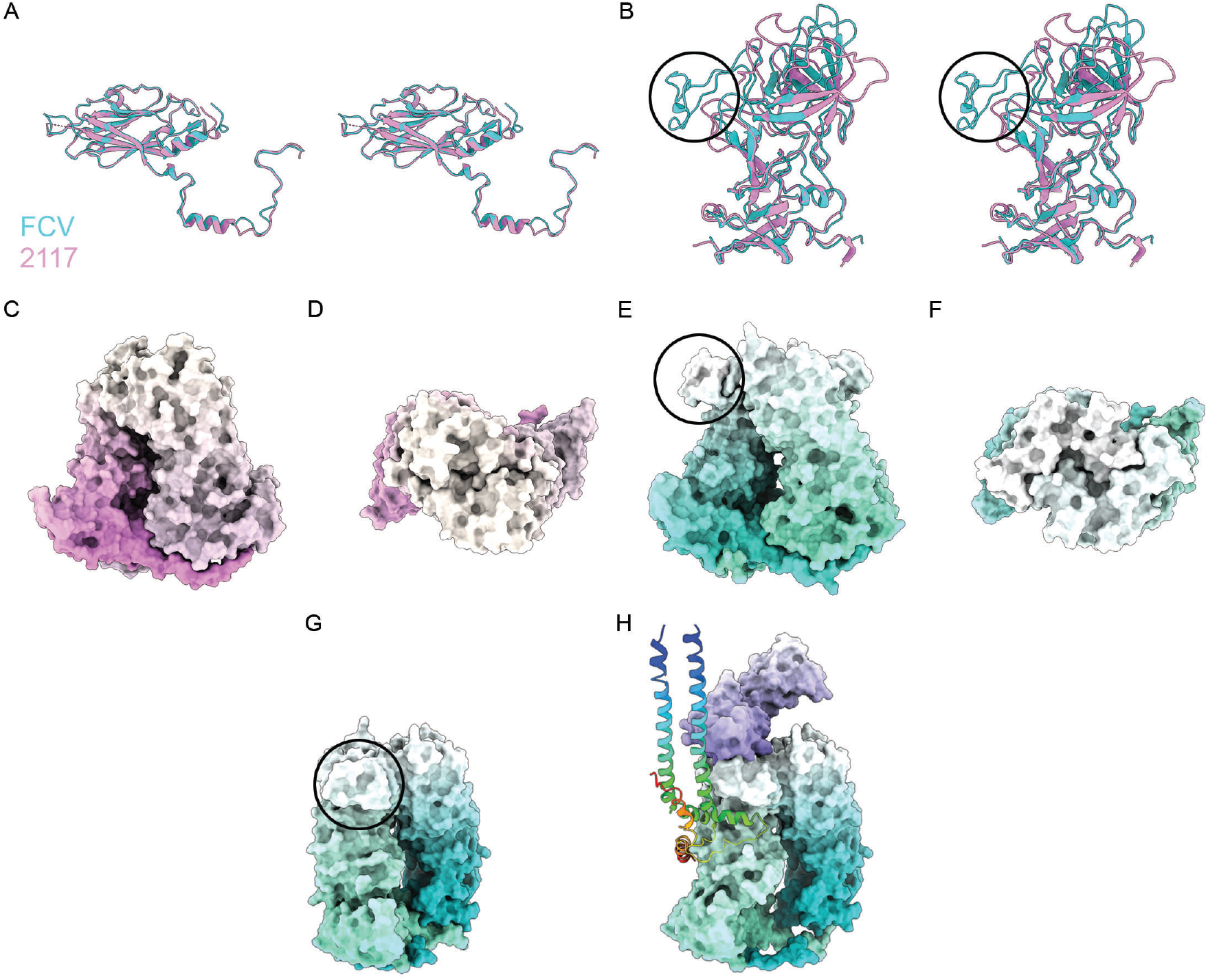
Comparison of the VP1 structure for vesivirus 2117 (violet) with that of FCV (PDB ID 6GSH - turquoise) shows a high degree of similarity in the S-domain (wall-eyed stereo pair view - A). The P-domains of 2117 and FCV show similar structure in the P1 sub-domains, whereas the P2 sub-domains exhibit major differences (wall-eyed stereo pair view - B). The most notable difference between the P2 regions of these two capsid proteins is the presence of an insert in the sequence of FCV VP1 that results in the formation of an extended loop or cantilevered arm (black circle B, E, G). Solvent excluded surface representations of the AB dimer of 2117 highlight the rounded shape of capsomeres in this virus, a consequence of the absence of the cantilevered arm. Side (C) and top (D) views are shown. Dimeric capsomeres of FCV VP1 form a flat platform on their outermost face as a consequence of this large insert, that appears rhombus shaped when viewed normal to the capsid surface (AB dimer shown, side – E, top – F). Receptor engagement leads to rearrangement of the cantilever feature, moving up towards the receptor (feline junctional adhesion molecule A – mauve), to form a cleft, into which VP2 binds – shown as a rainbow coloured ribbon diagram (G,H – PDB ID 6GSI).

### An insertion in the P2 domain of FCV gives rise to a functionally important extended loop that is absent from 2117 VP1

Of particular note, a major structural difference between P2 domains of 2117 and FCV is the presence of a large insertion (22 amino acid residues) in the FCV sequence, between β-strands two and three of the P2 beta-barrel. This region corresponds to amino-acid residues 409-468 in FCV VP1 and 296-333 in 2117 VP1. Sequence similarity in this region is low, thus defining a precise location for the insert is not possible. The additional sequence in this loop gives rise to the formation of a large cantilevered arm at amino-acid residues 430-463, conferring a rhomboid shape on the FCV P-dimer when viewed normal to the capsid floor (Fig 2B,E-F, Movie S2), while absence of this feature lends the 2117 P-dimer a more rounded shape (Fig 2C-D).

The cantilevered arm is of functional importance to FCV and its absence in 2117 points to a number of likely biological differences between the two Vesivirus clades. When FCV engages its cellular receptor - fJAM-A, the whole capsid undergoes substantial structural rearrangements. When saturated with soluble receptor fragments, we previously observed that P-dimers become more mobile, breaking icosahedral symmetry by rotating and tilting. At a unique three-fold symmetry axis, a pore opens in the capsid shell and a dodecameric funnel-shaped assembly of VP2 forms (15). We have previously proposed that VP2 is extruded from the capsid interior as we found no evidence of VP2 on capsids prior to receptor engagement, and that the function of this assembly is to act as a portal, mediating genome release across the endosomal membrane. To accommodate assembly of the VP2 portal the VP1 P domain undergoes several structural rearrangements, the most dramatic of which is lifting of the cantilevered arm towards the receptor. This results in widening of a cleft on the side of P2, into which a helix of VP2 inserts itself, anchoring the portal to the capsid surface (Fig 2G,H). VP2 is encoded by all caliciviruses and is known to be of critical importance (12), thus we expect 2117 VP2 to form a similar structure, however the mode of attachment to VP1 may not resemble that which we have previously described for FCV.

The distinct P2 topologies of 2117 and FCV point to substantial differences in receptor engagement as well as VP2 attachment. FCV P-dimers have a rhombus shaped outer face, providing an extended flattened platform onto which fJAM-A may bind (Fig 2H). When decorated with soluble fragments of the fJAM-A ectodomain, we previously observed disruption of the dimeric solution state of the receptor and possible formation of a new homodimerization interaction. The more rounded outer-face of 2117 P-dimers may accommodate a different mode of receptor engagement, perhaps at a more acute angle relative to the capsid surface. At the present time the cellular receptors for 2117 or other caliciviruses in the same clade are not known, however.

### Comparison of 2117 VP1 to other calicivirus capsid structures reveals a similarity to rabbit haemorrhagic disease virus

We have previously observed that the rounded shape of 2117 P-dimers more closely resembled the capsomeres of other caliciviruses, in particular Sapoviruses. Comparison of the 2117 P-domain from our atomic model to other known high-resolution structures for P indicate a lower degree of similarity to a Norovirus P, pairwise alignment and comparison of chain A of the 2117 P-domain to that of the snow-mountain strain of norovirus (PDB ID 6OTF (21)) indicated an RMSD value of 11.8 Å, while for murine norovirus (PDB ID 6IUK (22)) the difference was 13.8 Å. Interestingly the P-domain for rabbit haemorrhagic disease virus (RHDV - PDB ID 4X1W (23)), yielded an RMSD measurement of 5.2 Å in comparison with chain A of 2117 P, with very close agreement of the P1 domain in particular (Fig 3A). The S-domain of RHDV VP1 (PDB ID 4EJR (24)) also showed a highly similar structure; giving an RMSD of 2.3 Å when compared to chain C of 2117 VP1 (Fig 3B). No high-resolution data for a Sapovirus P are available at the time of writing, nonetheless the observed similarity to the major capsid protein of RHDV is intriguing.

**Figure 3.**
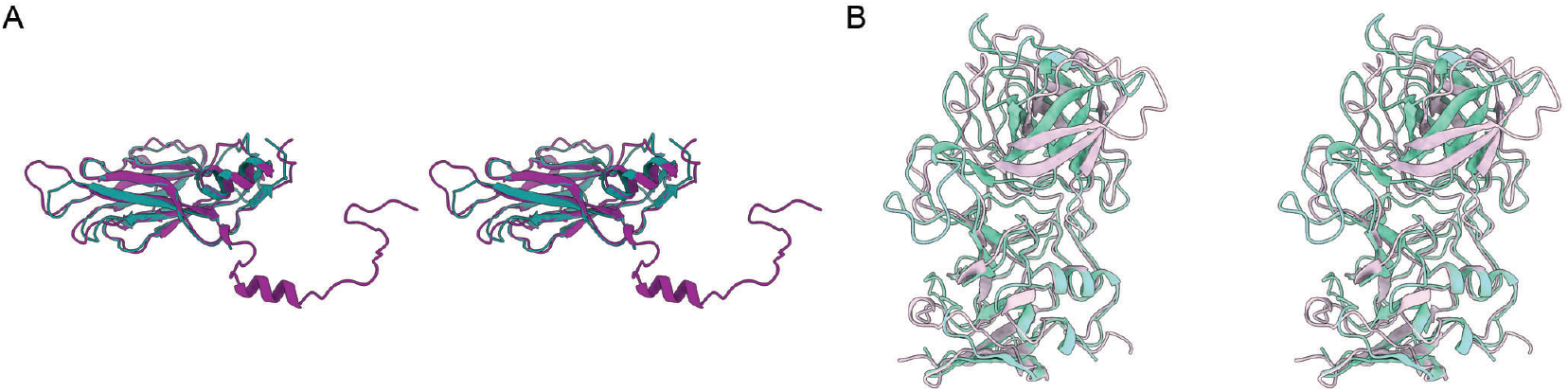
Vesivirus 2117 VP1 has a similar structure to the capsid protein of rabbit haemorrhagic disease virus. The X-ray structure for RHDV (PDB ID 4EJR) S-domain (teal) gave an RMSD of 2.3 Å, compared to the 2117 S-domain for chain C (dark-magenta, wall-eyed stereo view A). The X-ray structure for the P-domain of RHDV (PDB ID 4X1W – aquamarine), most closely resembled chain B for 2117 (thistle), showing a highly similar P1 sub-domain.

### Summary

We have determined the structure of vesivirus 2117 capsids by cryo-EM imaging of virus-like particles and computational three-dimensional image reconstruction. Our data allowed us to assemble an atomic model of the major capsid protein VP1, revealing several differences between this structure and the capsid of another Vesivirus - feline calicivirus, particularly in the P2 domain. The most note-worthy difference is absence of the cantilevered arm, a feature of the FCV P2 domain that rearranges following receptor engagement to accommodate the attachment of VP2. The lack of this loop in the vesivirus 2117 VP1 results in rounded capsomeres that more closely resemble those of other *Caliciviridae*, in particular the Lagovirus RHDV. Our findings suggest major functional differences in both receptor engagement and VP2 portal assembly between the FCV clade of the Vesiviruses and other *Caliciviridae*.

## Methods

### VLP production and cryo-electron microscopy

VLPs were prepared as previously described (6). Briefly the 2117 major capsid protein was expressed in Hi5 cells by baculovirus expression of the VP1 gene. Assembled particles were purified by differential ultracentrifugation and dialysed into phosphate buffered saline for imaging by cryo-EM. Five microlitres of purified VLPs were loaded onto freshly glow-discharged C-flat holey carbon support films (R2/2 ProtoChips) bearing a thin continuous carbon-film. VLPs were allowed to adhere to the carbon support for one minute before grids were blotted for 2 seconds and plunged into a bath of liquid nitrogen cooled liquid ethane. Specimen vitrification was performed in a Thermo-Fisher Vitrobot Mark IV held at 4°C and 100% humidity. Vitrified samples were imaged in a JEOL CryoARM 300 automated cryo transmission electron microscope equipped with a Direct Electron DE64 detector. Images were recorded at an accelerating voltage of 300 keV and magnification of 60k× which corresponds to a pixel size of 0.998 Å at the specimen scale. Two second exposures were captured at 25 frames per second with the detector operated in integrating mode and an electron flux of 1.1 electrons per Å^2^ per frame. Two thousand (2000) micrographs were recorded using the JADAS automation package.

### Image processing and three-dimensional, image reconstruction and model building

Image processing and three-dimensional image reconstruction was performed using Relion 3.0.3 (25, 26). Micrograph movies were motion corrected using MotionCor2 (27) and defocus estimation was accomplished using GCTF (28). An initial data set of 65071 particle images was selected for 2D classification followed by icosahedral 3D reconstruction and classification using our previously calculated 10 Å structure for vesivirus 2117 as a starting model. Following successive rounds of classification, a final dataset of 15989 particles was identified that yielded a refined 3D cryo-EM density map at 3.6 Å resolution according to the gold-standard criterion with an FSC cut off of 0.143. Local resolution assessment indicated that resolution ranged between 3.4 and 4.4 Å. Automated sharpening as implemented in Relion’s post-processing routine resulted in a map for which the NTA, S and P1 regions were well resolved but distal regions of P2 were poorly resolved and showed discontinuous densities. Similarly filtering the map according to local resolution did not yield interpretable P2 density. Deep learning map modification performed using deepEMhancer, yielded density that was able to be interpreted to build an atomic model. Retrospective comparison of the modified density to a map that had been blurred with a B-factor of 50 Å-1 using MRC to MTZ within the CCP-EM package, alongside the presence of quasi-symmetry in the icosahedral reconstruction confirmed that map modification was appropriate (20, 29–31). A starting model was generated by submitting the protein sequence for 2117 VP1 to the Phyre-2 server (19). The homology model coordinates were docked to the cryo-EM density map using UCSF Chimera (32) and manually edited using Coot v0.9.2 (33). Real-space and reciprocal space refinement of coordinates was performed using PHENIX (34) and REFMAC (30) respectively. Severe clashes and rotamer outliers were corrected using ISOLDE implemented in UCSF ChimeraX (35, 36). ChimeraX was also used for all visualisation and interpretation tasks. Validation of coordinates by the protein data-bank server indicated a clashscore of 5, Ramachandran outliers of 0.5% and sidechain outliers of 0.5%. Phenix validation indicated a MolProbity score of 1.9.

## Acknowledgements

We acknowledge the Scottish Centre for Macromolecular Imaging (SCMI) for access to cryo-EM instrumentation, funded by the Medical Research Council (MC_ PC_17135) and Scottish Funding Council (H17007). DB and JS are funded by the Medical Research Council (MC_ UU_12014/7), MJC is funded by the Biotechnology and Biological Sciences Research Council (BB/T002239/1). IG is a Wellcome Senior Fellow and supported by funding from the Wellcome Trust (REF: 207498/Z/17/Z).

The MRC-University of Glasgow Centre for Virus Research uses the CRediT taxonomy of author contributions. HS – investigation, formal analysis, writing – original draft. MJC – formal analysis, funding acquisition, supervision, writing - review and editing. EE – resources, writing – review and editing. JS – investigation, writing – review and editing. IGG – conceptualisation, funding acquisition, resources, writing – review and editing. DB – concept0075alisation, data curation, investigation, formal analysis, funding acquisition, supervision, validation, visualisation, writing – original draft.

## Data Deposition

Cryo-EM maps have been deposited in the EM-databank (https://www.ebi.ac.uk/pdbe/emdb/) with the accession number EMD-12194.

Coordinates have been deposited in the protein data bank (https://www.ebi.ac.uk/pdbe/) with the accession number 7BJP

The raw micrograph movies are deposited in the EMPI-AR databank (https://www.ebi.ac.uk/pdbe/emdb/empiar/) with accession number ******.

## References

1. Oehmig A, Buttner M, Weiland F, Werz W, Bergemann K, Pfaff E. 2003. Identification of a calicivirus isolate of unknown origin. J Gen Virol 84:2837–2845.

2. Qiu Y, Jones N, Busch M, Pan P, Keegan J, Zhou W, Plavsic M, Hayes M, McPherson JM, Edmunds T, Zhang K, Mattaliano RJ. 2013. Identification and quantitation of Vesivirus 2117 particles in bioreactor fluids from infected Chinese hamster ovary cell cultures. Biotechnol Bioeng 110:1342–53.

3. Vinje J, Estes MK, Esteves P, Green KY, Katayama K, Knowles NJ, L’Homme Y, Martella V, Vennema H, White PA, Ictv Report C. 2019. ICTV Virus Taxonomy Profile: *Caliciviridae*. J Gen Virol 100:1469–1470.

4. Kapikian AZ, Wyatt RG, Dolin R, Thornhill TS, Kalica AR, Chanock RM. 1972. Visualization by immune electron microscopy of a 27-nm particle associated with acute infectious nonbacterial gastroenteritis. J Virol 10:1075–81.

5. Pedersen NC, Elliott JB, Glasgow A, Poland A, Keel K. 2000. An isolated epizootic of hemorrhagic-like fever in cats caused by a novel and highly virulent strain of feline calicivirus. Vet Microbiol 73:281–300.

6. Conley M, Emmott E, Orton R, Taylor D, Carneiro DG, Murata K, Goodfellow IG, Hansman GS, Bhella D. 2017. Vesivirus 2117 capsids more closely resemble sapovirus and lagovirus particles than other known vesivirus structures. J Gen Virol 98:68–76.

7. Renshaw RW, Griffing J, Weisman J, Crofton LM, Laverack MA, Poston RP, Duhamel GE, Dubovi EJ. 2018. Characterization of a Vesivirus Associated with an Outbreak of Acute Hemorrhagic Gastroenteritis in Domestic Dogs. J Clin Microbiol 56.

8. Di Martino B, Di Profio F, Lanave G, De Grazia S, Giammanco GM, Lavazza A, Buonavoglia C, Marsilio F, Banyai K, Martella V. 2015. Antibodies for strain 2117-like vesiviruses (caliciviruses) in humans. Virus Res 210:279–82.

9. Di Martino B, Di Profio F, Melegari I, Sarchese V, Massirio I, Luciani A, Lanave G, Marsilio F, Martella V. 2018. Serological and molecular investigation of 2117-like vesiviruses in cats. Arch Virol 163:197–201.

10. Neill JD. 1990. Nucleotide sequence of a region of the feline calicivirus genome which encodes picornavirus-like RNA-dependent RNA polymerase, cysteine protease and 2C polypeptides. Virus Res 17:145–60.

11. Neill JD, Reardon IM, Heinrikson RL. 1991. Nucleotide sequence and expression of the capsid protein gene of feline calicivirus. J Virol 65:5440–7.

12. Sosnovtsev SV, Belliot G, Chang KO, Onwudiwe O, Green KY. 2005. Feline calicivirus VP2 is essential for the production of infectious virions. J Virol 79:4012–24.

13. Tohya Y, Taniguchi Y, Takahashi E, Utagawa E, Takeda N, Miyamura K, Yamazaki S, Mikami T. 1991. Sequence analysis of the 3’-end of feline calicivirus genome. Virology 183:810–4.

14. Chen R, Neill JD, Estes MK, Prasad BV. 2006. X-ray structure of a native calicivirus: structural insights into antigenic diversity and host specificity. Proc Natl Acad Sci U S A 103:8048–53.

15. Conley MJ, McElwee M, Azmi L, Gabrielsen M, Byron O, Goodfellow IG, Bhella D. 2019. Calicivirus VP2 forms a portal-like assembly following receptor engagement. Nature 565:377–381.

16. Makino A, Shimojima M, Miyazawa T, Kato K, Tohya Y, Akashi H. 2006. Junctional adhesion molecule 1 is a functional receptor for feline calicivirus. J Virol 80:4482–90.

17. Bhella D, Gatherer D, Chaudhry Y, Pink R, Goodfellow IG. 2008. Structural insights into calicivirus attachment and uncoating. J Virol 82:8051–8.

18. Bhella D, Goodfellow IG. 2011. The cryo-electron microscopy structure of feline calicivirus bound to junctional adhesion molecule A at 9-angstrom resolution reveals receptor-induced flexibility and two distinct conformational changes in the capsid protein VP1. J Virol 85:11381–90.

19. Kelley LA, Sternberg MJ. 2009. Protein structure prediction on the Web: a case study using the Phyre server. Nat Protoc 4:363–71.

20. Sanchez-Garcia R, Gomez-Blanco J, Cuervo A, Carazo J, Sorzano C, Vargas J. 2020. DeepEMhancer: a deep learning solution for cryo-EM volume post-processing. bioRxiv.

21. Jung J, Grant T, Thomas DR, Diehnelt CW, Grigorieff N, Joshua-Tor L. 2019. High-resolution cryo-EM structures of outbreak strain human norovirus shells reveal size variations. Proc Natl Acad Sci U S A 116:12828–12832.

22. Song C, Takai-Todaka R, Miki M, Haga K, Fujimoto A, Ishiyama R, Oikawa K, Yokoyama M, Miyazaki N, Iwasaki K, Murakami K, Katayama K, Murata K. 2020. Dynamic rotation of the protruding domain enhances the infectivity of norovirus. PLoS Pathog 16:e1008619.

23. Leuthold MM, Dalton KP, Hansman GS. 2015. Structural analysis of a rabbit hemorrhagic disease virus binding to histo-blood group antigens. J Virol 89:2378–87.

24. Wang X, Xu F, Liu J, Gao B, Liu Y, Zhai Y, Ma J, Zhang K, Baker TS, Schulten K, Zheng D, Pang H, Sun F. 2013. Atomic model of rabbit hemorrhagic disease virus by cryo-electron microscopy and crystallography. PLoS Pathog 9:e1003132.

25. Scheres SH. 2012. RELION: implementation of a Bayesian approach to cryo-EM structure determination. J Struct Biol 180:519–30.

26. Zivanov J, Nakane T, Forsberg BO, Kimanius D, Hagen WJ, Lindahl E, Scheres SH. 2018. New tools for automated high-resolution cryo-EM structure determination in RELION-3. Elife 7.

27. Zheng SQ, Palovcak E, Armache JP, Verba KA, Cheng Y, Agard DA. 2017. MotionCor2: anisotropic correction of beam-induced motion for improved cryo-electron microscopy. Nat Methods 14:331–332.

28. Zhang K. 2016. Gctf: Real-time CTF determination and correction. J Struct Biol 193:1–12.

29. Burnley T, Palmer CM, Winn M. 2017. Recent developments in the CCP-EM software suite. Acta Crystallogr D Struct Biol 73:469–477.

30. Murshudov GN, Skubak P, Lebedev AA, Pannu NS, Steiner RA, Nicholls RA, Winn MD, Long F, Vagin AA. 2011. REFMAC5 for the refinement of macromolecular crystal structures. Acta Crystallogr D Biol Crystallogr 67:355–67.

31. Nicholls RA, Tykac M, Kovalevskiy O, Murshudov GN. 2018. Current approaches for the fitting and refinement of atomic models into cryo-EM maps using CCP-EM. Acta Crystallogr D Struct Biol 74:492–505.

32. Pettersen EF, Goddard TD, Huang CC, Couch GS, Greenblatt DM, Meng EC, Ferrin TE. 2004. UCSF Chimera--a visualization system for exploratory research and analysis. J Comput Chem 25:1605–12.

33. Emsley P, Lohkamp B, Scott WG, Cowtan K. 2010. Features and development of Coot. Acta Crystallogr D Biol Crystallogr 66:486–501.

34. Liebschner D, Afonine PV, Baker ML, Bunkoczi G, Chen VB, Croll TI, Hintze B, Hung LW, Jain S, McCoy AJ, Moriarty NW, Oeffner RD, Poon BK, Prisant MG, Read RJ, Richardson JS, Richardson DC, Sammito MD, Sobolev OV, Stockwell DH, Terwilliger TC, Urzhumtsev AG, Videau LL, Williams CJ, Adams PD. 2019. Macromolecular structure determination using X-rays, neutrons and electrons: recent developments in Phenix. Acta Crystallogr D Struct Biol 75:861–877.

35. Croll TI. 2018. ISOLDE: a physically realistic environment for model building into low-resolution electron-density maps. Acta Crystallogr D Struct Biol 74:519–530.

36. Pettersen EF, Goddard TD, Huang CC, Meng EC, Couch GS, Croll TI, Morris JH, Ferrin TE. 2021. UCSF ChimeraX: Structure visualization for researchers, educators, and developers. Protein Sci 30:70–82.

