## Supplemental figures for "The cryo-EM structure of vesivirus 2117 highlights functional variations in entry pathways for viruses in different clades of the Vesivirus genus"

Supplemental data

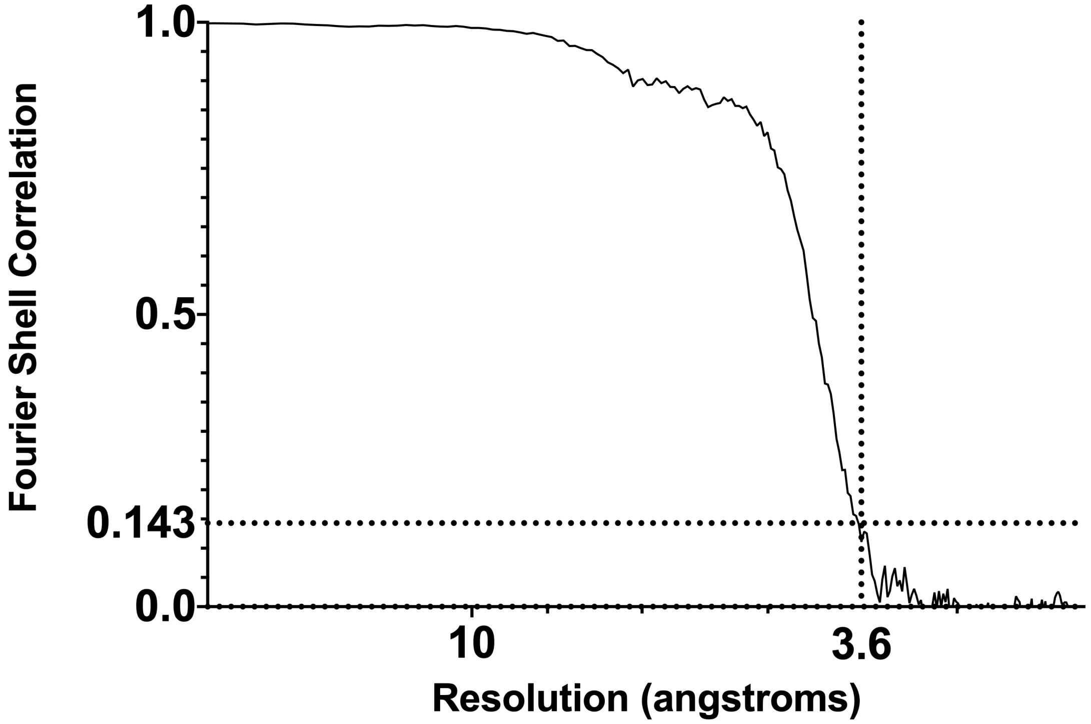


***Figure S1***

Fourier shell correlation plot for the cryo-EM reconstruction of vesivirus 2117 virus-like particles. Half-maps were calculated according to the gold-standard method. Correlation crosses the 0.143 cut-off at a resolution of 3.6 angstroms.

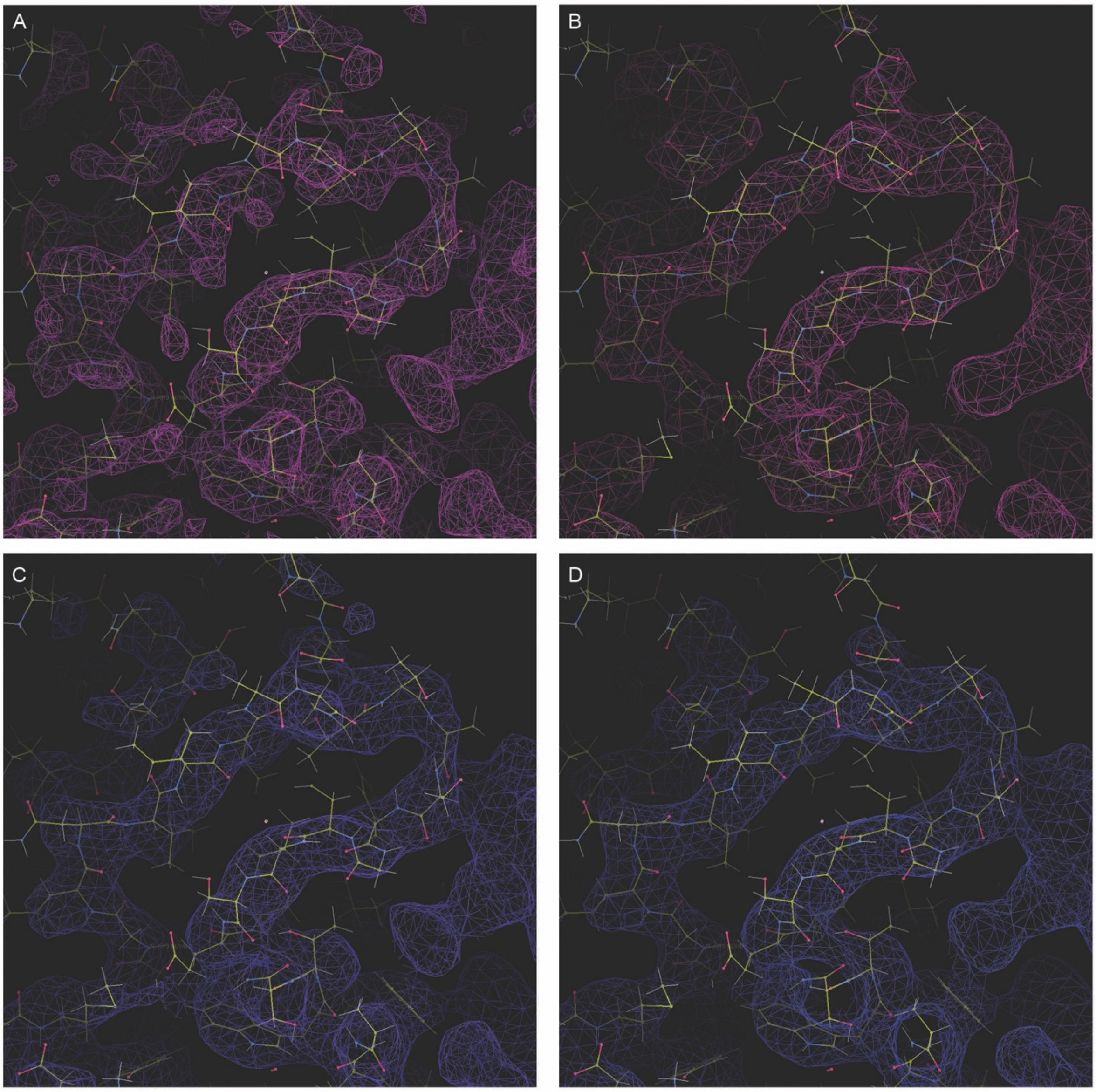


***Figure S2***
Comparison of cryo-EM maps for amino-acid residue 315-330 in chain B, a region where the sharpened map presented noisy discontinuous density (A). DeepEMhancer modification facilitated tracing of the polypeptide chain in ambiguous and noisy density (B). A similar level of improvement was subsequently found to be possible by blurring maps; applying a B-factor of 50 Å $^{-1}$ (C) or 100 Å $^{-1}$ (D).

**Movie Descriptions.**
Time points are given for each described event

***Movie S1***
Movie to show three-dimensional reconstructions of the vesivirus 2117 virus-like particle and resulting atomic models. (9s) Isosurface representation of the sharpened cryo-EM density map coloured according to radius. This map reveals noisy and indistinct density in the distal regions of capsomeres. (36s) Local resolution assessment indicates a resolution of poorer than 4 angstroms for this region, which in other caliciviruses incorporates receptor binding sites and major immunodominant epitopes. (37s) Deep-learning map modification was applied to enhance density, allowing model building to proceed.(58s) The asymmetric unit of the T=3 icosahedral capsid comprises three copies of VP1, A (violet), B (thistle) and C (deep magenta). (1m16s) An atomic model was built into the modified cryo-EM density and is shown as a ribbon diagram of the asymmetric unit (1m24s) and AB dimer (1m56s) and CC dimer (2m37s).

***Movie S2***
Movie to show structural comparison of 2117 VP1 to FCV VP1. The asymmetric unit of the 2117 capsid is shown as a ribbon diagram (8s). Comparison of the S domain reveals a highly conserved structure (39s). Comparison of the P domains shows that while P1 is similar, the P2 sub-domain has major differences in surface topology (1m10s). The most notable difference is the presence of a 22 amino-acid residue insertion on the sequence of FCV; a large extended loop forming a cantilevered arm (1m36s). A solvent excluded surface representation of the 2117 AB dimer highlights the rounded capsomere shape (1m59s). The FCV AB-dimer however presents a rhombus shaped outer face as a consequence of the presence of the cantilevered arm (2m29s). Following receptor engagement, this feature moves upwards to engage the fJAM-A molecule leading to the opening of a cleft on the side of P2, into which VP2 binds (3m06s).
